# Task-evoked functional connectivity exhibits novel and strengthened relationships with executive function relative to the resting state

**DOI:** 10.1101/2025.06.30.661993

**Authors:** Mackenzie E. Mitchell, Eric Feczko, Damien A. Fair, Jessica R. Cohen

## Abstract

Executive functioning in children has been linked to intrinsic brain network organization assessed during the resting state, as well as to brain network organization during the performance of cognitive tasks. Prior work has established that task-based brain networks are stronger predictors of behavior than resting state networks, yet it is unclear if tasks only strengthen relationships that exist weakly at rest or if tasks also evoke unique relationships. A lack of discernment regarding how tasks and the resting state commonly and uniquely support executive functions precludes a holistic understanding of the neurobiological basis of executive functions. This project investigated differences in brain network organization and relationships with executive function ability between the resting state and two executive function tasks, a stop signal task and an emotional n-back task, using the Adolescent Brain and Cognitive Development (ABCD) Study dataset. Both executive function tasks evoked a more integrated network organization than the resting state, and executive function ability was related to different aspects of brain network organization during the resting state and during the tasks. Further, task-related shifts in brain network organization evoked several new relationships with executive function that were not detectable during the resting state and strengthened a relationship with executive function that existed weakly during the resting state. Overall, this study establishes a distinction between common and unique features of intrinsic and task-evoked brain function that facilitate executive function in children.

**Significance:** Executive functions, which encompass goal-directed behaviors critical for life success, emerge from interactions within and between networks of brain regions. Here, we tested how executive functions are linked to functional brain network interactions during the resting state, in which there are no external cognitive demands, and during executive function tasks in late childhood. We found that brain network organization during the performance of executive function tasks evoked a combination of new and strengthened relationships with executive function ability compared to the resting state. This suggests that the brain facilitates executive function performance in children through the recruitment of specific functional interactions during the enactment of executive function.

## Introduction

Response inhibition and working memory are core domains of executive function (Diamond, 2013; Miyake et al., 2000) that emerge early in life and develop slowly throughout childhood and adolescence (Best & Miller, 2010; Tervo-Clemmens et al., 2023). A large body of research has linked patterns of functional connectivity during the resting state to executive function ability in both youth and adults (Avery et al., 2020; Gu et al., 2015, 2020; Keller et al., 2023; Marek et al., 2019; Reineberg et al., 2015; Reineberg & Banich, 2016; Satterthwaite et al., 2015; Tooley et al., 2022). These studies emphasize that the maturation of executive function is supported by the modular development of intrinsic functional brain networks (Gu et al., 2015; Keller et al., 2023; Tooley et al., 2022), or networks of strongly interconnected regions that exhibit relatively sparse connectivity across networks. Examining functional connectivity during rest is useful for understanding how executive function and other cognitive processes emerge because resting state, or intrinsic, functional connectivity is thought to reflect historical coactivation of brain regions (Dosenbach et al., 2007; Kelly et al., 2008) and homeostatic processes (Laumann & Snyder, 2021).

Yet examining functional connectivity while completing executive function tasks may more directly capture on-line cognitive processing (Finn, 2021). Empirical literature has robustly demonstrated small but meaningful differences in functional brain network organization between the resting state and cognitive tasks in both adults (Cohen & D’Esposito, 2016; Cole et al., 2014; Gratton, Laumann, et al., 2018; Krienen et al., 2014; Schultz & Cole, 2016; Shine et al., 2016) and children (Dwyer et al., 2014; Le et al., 2020; Mitchell et al., 2024). Tasks of executive function, including working memory, generally demonstrate significantly greater integration across the whole brain relative to the resting state in adults (Braun et al., 2015; Cohen & D’Esposito, 2016; Shine et al., 2016) and children (Le et al., 2020), as well as greater integration between particular networks involved in cognitive control and attention (Kolskår et al., 2018). These differences in network organization between intrinsic and task-based states probing executive function may indicate the presence of unique on-line neurocognitive mechanisms undergirding executive function. In line with this, several recent studies have reported that task-based functional connectivity from a variety of tasks is more predictive of behavior than resting state functional connectivity in both adults (Greene et al., 2018, 2020; Jiang et al., 2020) and children (J. Chen et al., 2022; Zhao et al., 2023). Additionally, studies utilizing task-based functional connectivity in children and adolescents identify relationships with executive function that are not observed in the resting state literature. For example, better response inhibition in youth is related to greater segregation, or anticorrelations, between regions of cognitive control-related networks and regions of the default mode network during tasks (Dwyer et al., 2014; Stevens et al., 2007), but this relationship is not present during the resting state in children (Barber et al., 2013; Mennes et al., 2012). Relationships with working memory similarly show differences between task-based and resting state functional connectivity, with evidence that better working memory in children is related to more segregation during the resting state (Zhong et al., 2014) but more integration during a working memory task (M. Chen et al., 2022). Therefore, understanding how executive functions are facilitated by both intrinsic brain organization during the resting state and task-evoked brain organization will enable a more comprehensive understanding of the neurobiological basis of executive function.

The distinction between resting state and task-based facilitation of executive function ability is important for understanding the neurobiological mechanisms that give rise to individual differences in performance on executive function tasks. Building off the work from Greene and colleagues (2018, 2020), one possible explanation for differences in executive function ability could be that during executive function tasks individuals engage executive-function specific network substrates to varying degrees. An additional possibility is that individual differences in intrinsic brain network organization impact the degree of network reconfiguration needed to successfully meet executive function demands. This idea is supported by evidence that better task performance in adults is related to a higher degree of similarity in patterns of functional connectivity between the resting state and tasks, suggesting that task-efficient organization is reliant upon optimized resting state organization (Schultz & Cole, 2016). Therefore, we hypothesize that executive function is supported by a combination of brain network organization features that are present during both the resting state and tasks, as well as unique features specific to each state.

In this project, we tested for differences in functional brain organization between the resting state and two task-evoked executive function states using baseline data from the ABCD Study when participants were 9-10 years old. We looked at relationships between brain network metrics and performance on a stop signal task (SST) probing response inhibition and an emotional n-back (EN-back) task probing working memory. Of particular interest, we tested for differences in relationships with task performance between the resting state and task-evoked states. Understanding how resting state and task-evoked functional connectivity relate to response inhibition and working memory task performance in childhood will serve to distinguish how executive functions are supported by both intrinsic processes and on-line cognitive strategies during tasks.

## Results

Here, we examined differences in large-scale brain network organization and relationships with executive function ability between the resting state and two executive function tasks: the SST and EN-back. We used two large, demographically matched samples (discovery: *N* = 2,470, replication: *N* = 2,479) from the ABCD Study (Casey et al., 2018; Feczko et al., 2020) to ensure reliability of our results (**Table 1**). We only report and discuss results that were significant in both the discovery and replication samples. See the **Supplement** for all results from both samples.

**Table 1.**
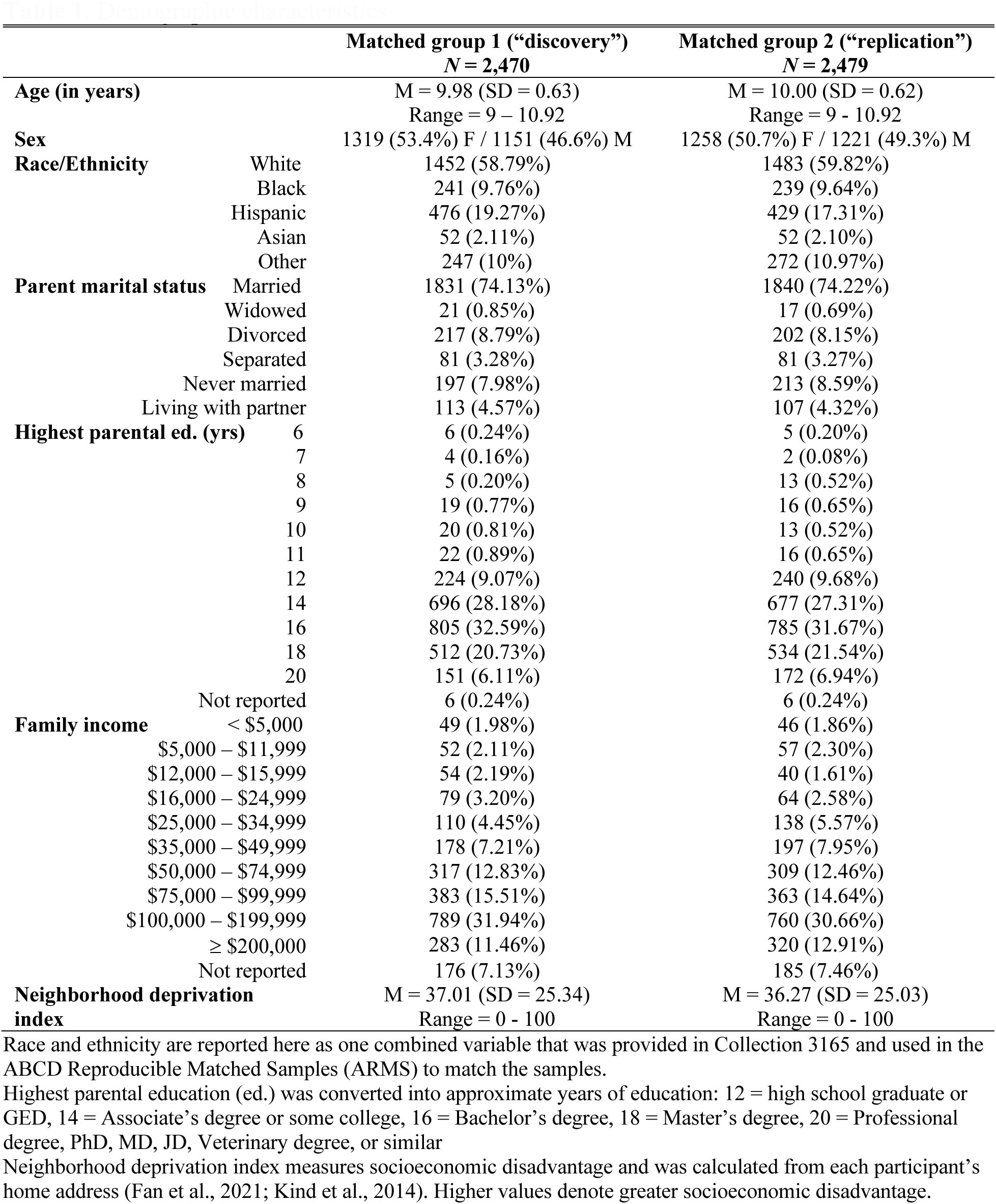
Demographic characteristics.

### Differences in brain network organization between the resting state and task-evoked states

To quantify differences in whole brain organization between the resting state and task-evoked states, modularity and global efficiency were estimated as measures of segregation and integration respectively. To quantify differences in integration of specific networks, network dissociation index was estimated as the average node dissociation index across all nodes within a given network.

During the SST as compared to the resting state, we observed significantly decreased modularity (discovery: ***β*** = −1.22, adjusted *p* < .001; replication: ***β*** = −1.10, adjusted *p* < .001), increased global efficiency (discovery: ***β*** = 0.12, adjusted *p* < .001; replication: ***β*** = 0.12, adjusted *p* < .001), and increased network dissociation index in all networks except for the fronto-parietal network (all significant adjusted *p*-values < .001; **Supplementary Table 1**).

During the EN-back task as compared to the resting state, we similarly observed significantly decreased modularity (discovery: ***β*** = −2.64, adjusted *p* < .001; replication: ***β*** = - 2.59, adjusted *p* < .001), increased global efficiency (discovery: ***β*** = 0.90, adjusted *p* < .001; replication: ***β*** = 0.90, adjusted *p* < .001), and increased network dissociation index in all networks (all adjusted *p*-values < .001; **Supplementary Table 2**).

### Relationships between brain network organization and task performance

Next, we quantified relationships between brain network organization and task performance. First, we assessed whether brain organization during the resting state or the SST were related to stop signal reaction time (SSRT) on the SST. There were no significant relationships between whole brain or network-level measures of resting state brain organization and the SSRT after multiple comparison correction (all adjusted *p*-values > .145). However, during the SST, decreased cingulo-parietal network dissociation index was related to increased SSRT, or worse task performance (discovery: ***β*** = −0.06, adjusted *p* = .018; replication: ***β*** = - 0.06, adjusted *p* = .027; all other adjusted *p*-values > .122; see **Figure 1a** and **Supplementary Table 3)**.

**Figure 1.**
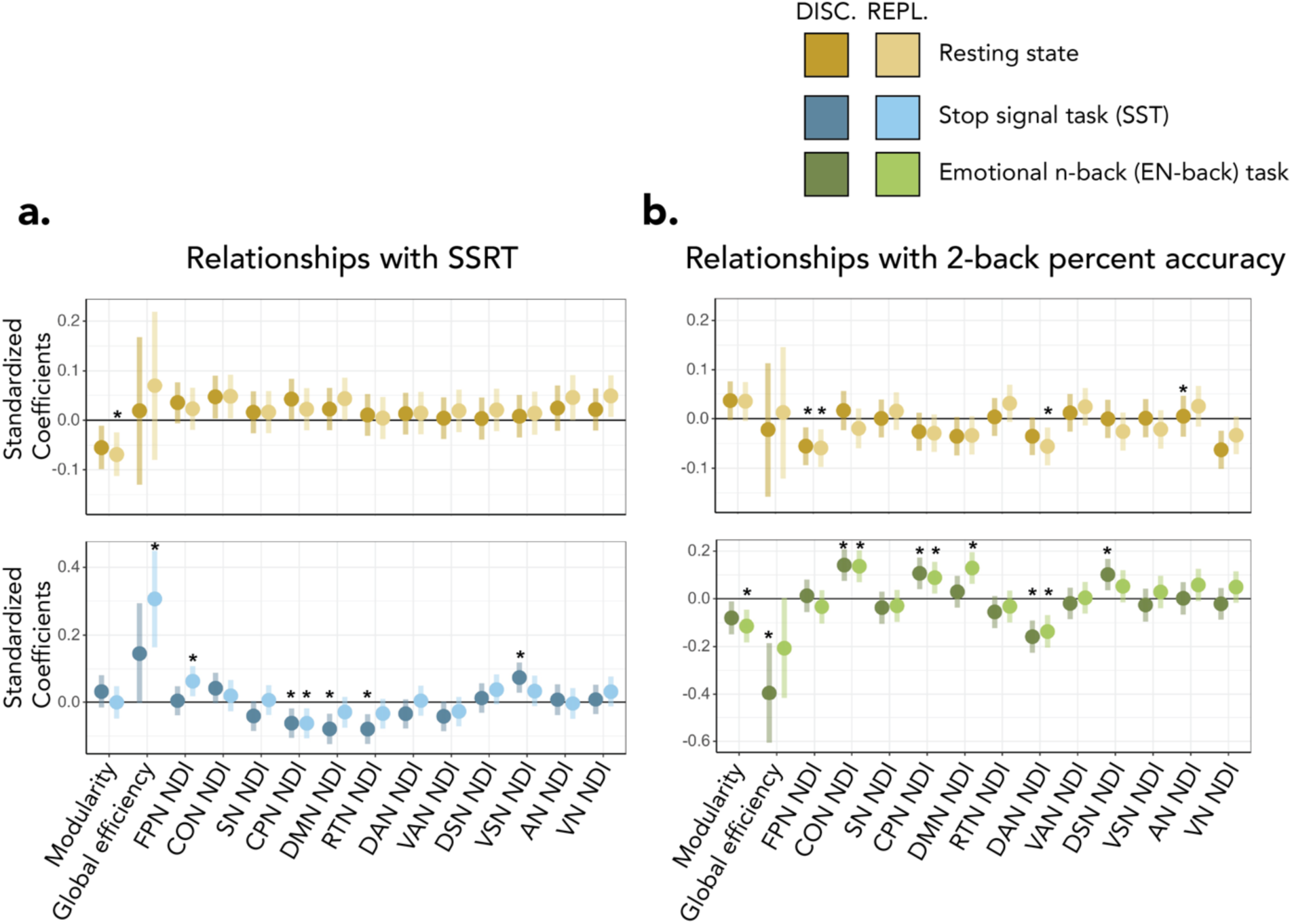
Relationships between resting state and task state brain metrics and performance on the **(a)** stop signal task and **(b)** emotional n-back task for both the discovery and replication samples. Significance after multiple comparison correction is denoted with asterisks: \**p*<.05. Y-axis limits are adjusted for each plot to display the error bars. DISC = discovery sample; REPL = replication sample; SSRT = stop signal reaction time; NDI = network dissociation index; FPN = fronto-parietal network; CON = cingulo-opercular network; SN = salience network; CPN = cingulo-parietal network; DMN = default mode network; RTN = retrosplenial-temporal network; DAN = dorsal attention network; VAN = ventral attention network; DSN = dorsal somatomotor network; VSN = ventral somatomotor network; AN = auditory network; VN = visual network

Then, we assessed whether brain network organization during the resting state or the EN-back task were related to 2-back accuracy on the EN-back task. During the resting state, decreased fronto-parietal network dissociation index was related to increased 2-back accuracy (discovery: ***β*** = −0.06, adjusted *p* = .029; replication: ***β*** = −0.06, adjusted *p* = .028). During the EN-back task, increased cingulo-opercular and cingulo-parietal network dissociation index, and decreased dorsal attention network dissociation index, were related to increased 2-back accuracy (cingulo-opercular: discovery: ***β*** = 0.14, adjusted *p* < .001; replication: ***β*** = 0.09, adjusted *p* = .027; cingulo-parietal: discovery: ***β*** = 0.11, adjusted *p* = .006; replication: ***β*** = 0.13, adjusted *p* = .001; dorsal attention: discovery: ***β*** = −0.16, adjusted *p* < .001; replication: ***β*** = −0.14, adjusted *p* = .001; all other adjusted *p*-values > .124; see **Figure 1b** and **Supplementary Table 4**).

### Modulation of brain-behavior relationships between the resting state and task states

Finally, we investigated whether relationships between functional brain network organization and task performance differed between the resting state and the task-evoked states. We focused on the significant brain-behavior relationships reported above. First, we compared brain-behavior relationships between the resting state and the SST. We found that cingulo-parietal network dissociation index had a significantly different relationship with SSRT during the resting state as compared to during the SST (discovery: ***β*** = −0.0018, adjusted *p* < .001; replication: ***β*** = −0.0015, adjusted *p* < .001). Specifically, cingulo-parietal network dissociation index was not significantly related to SSRT during the resting state but was significantly related to SSRT during the SST (**Figure 2**).

**Figure 2.**
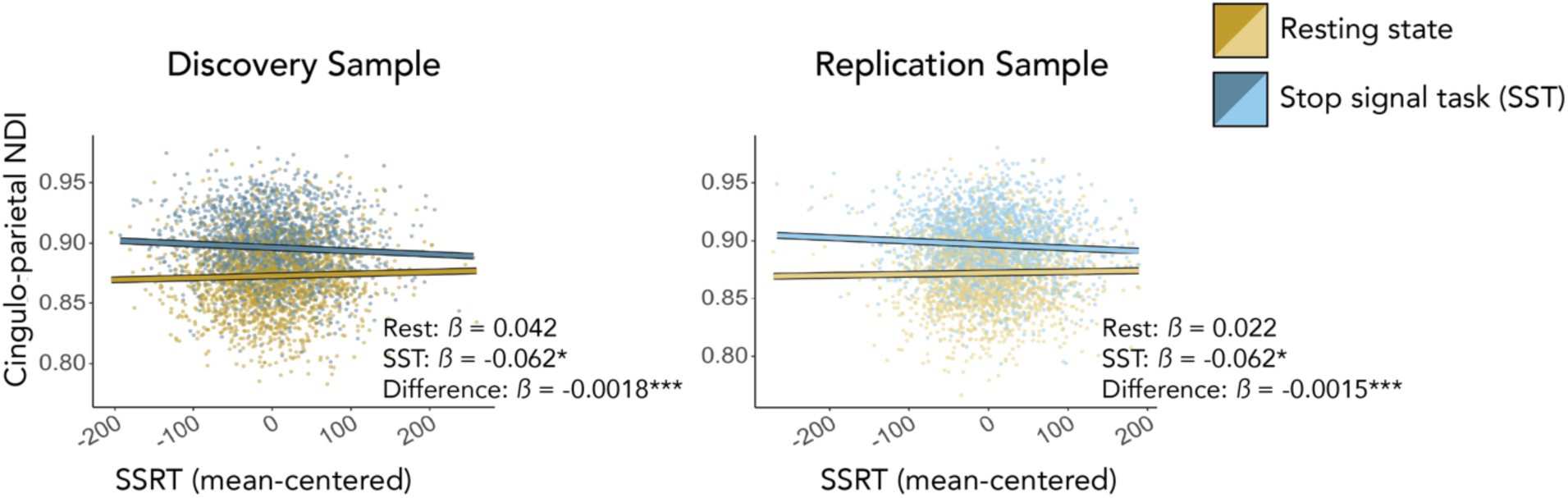
The relationship between cingulo-parietal network dissociation index and SSRT differs between the SST and the resting state. Relationships are shown for both the discovery (left) and replication (right) samples. Across both samples, cingulo-parietal network dissociation index during the resting state was not significantly related to SSRT. However, cingulo-parietal network dissociation index during the SST was significantly related to SSRT across both samples. The difference between the resting state and task-based relationships was significant in both samples. Standardized betas are reported. Significance after multiple comparison correction is denoted with asterisks: \**p*<.05, \*\*\**p*<.001. SSRT = stop signal reaction time; NDI = network dissociation index.

Next, we compared brain-behavior relationships between the resting state and the EN-back task. We found that although fronto-parietal network dissociation index had a significantly different relationship with 2-back accuracy across the resting state and the EN-back task for the discovery sample (discovery: ***β*** = 0.89, adjusted *p* = .044), this relationship was not consistent for the replication sample (replication: ***β*** = 0.11, adjusted *p* = .803). Even though the relationship was not significantly different across the resting state and the EN-back task for the replication sample, in both samples fronto-parietal network dissociation index was significantly related to 2-back accuracy during the resting state but not significantly related to 2-back accuracy during the EN-back task. The remainder of the significant results were consistent across the discovery and replication samples. We found significantly different relationships with 2-back accuracy during the resting state as compared to during the EN-back task for network dissociation index of the cingulo-opercular cingulo-parietal, and dorsal attention networks (cingulo-opercular: discovery: ***β*** = 1.71, adjusted *p* < .001; replication: ***β*** = 2.47, adjusted *p* < .001; cingulo-parietal: discovery: ***β*** = 1.36, adjusted *p* = .001; replication: ***β*** = 1.30, adjusted *p* < .001; dorsal attention: discovery: ***β*** = −2.11, adjusted *p* < .001; replication: ***β*** = −1.58, adjusted *p* < .001; **Figure 3**). Cingulo-opercular and cingulo-parietal network dissociation index were not significantly related to 2-back accuracy during the resting state but were significantly related to 2-back accuracy during the EN-back task. Dorsal attention network dissociation index was significantly related to 2-back accuracy during the resting state for the replication sample, but not for the discovery sample. However, during the EN-back task dorsal attention network dissociation index was more strongly significantly related to 2-back accuracy for both the discovery and replication samples. See **Supplementary Table 5**.

**Figure 3.**
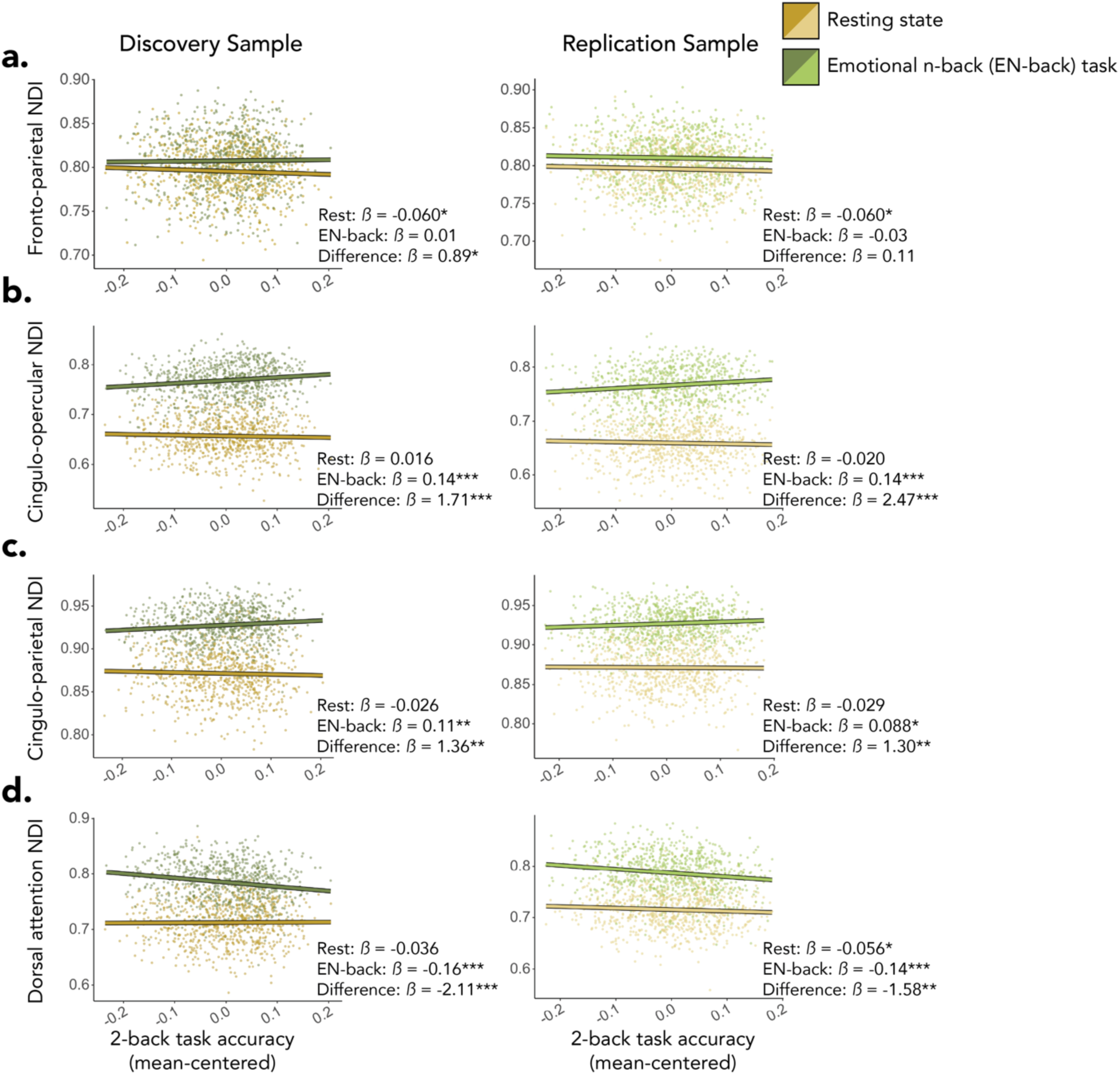
Relationships between network dissociation index and 2-back accuracy differ between the EN-back and the resting state. Relationships are shown for both the discovery (left) and replication (right) samples. **(a)** Across both samples, fronto-parietal network dissociation index during the resting state was significantly related to 2-back accuracy, while fronto-parietal network dissociation index during the EN-back task was not related to 2-back accuracy. The difference between the resting state and task-based relationships was significant in the discovery sample, but not in the replication sample. **(b-c)** Across both samples, cingulo-opercular and cingulo-parietal network dissociation index during the resting state were not related to 2-back accuracy. However, cingulo-opercular and cingulo-parietal network dissociation index during the EN-back task were significantly related to 2-back accuracy across both samples. Additionally, the difference between the resting state and task-based relationships was significant in both samples for both networks. **(d)** Dorsal attention network dissociation index during the resting state was related to 2-back accuracy in the replication, but not the discovery sample. However, dorsal attention network dissociation index during the EN-back task was significantly related to 2-back accuracy across both samples. Additionally, the difference between the resting state and task-based relationships was significant in both samples. Standardized betas are reported. Significance after multiple comparison correction is denoted with asterisks: \**p*<.05, \*\**p*<.01, \*\*\**p*<.001. NDI = network dissociation index.

## Discussion

In this study, we leveraged the ABCD Study baseline sample to investigate differences in functional brain network organization between the resting state and two tasks probing distinct aspects of executive function, a stop signal task (SST) and an emotional N-back (EN-back) task, in children aged 9-10 years. We assessed how brain network organization during the resting state and the executive function tasks related to executive function task performance, as well as whether the resting state and tasks elicited different relationships between functional brain organization and task performance. Our findings revealed lower modularity, higher global efficiency, and higher network dissociation index across most networks during both the SST and the EN-back task compared to the resting state, indicating decreased network segregation and increased network integration. Additionally, relationships with executive function task performance were found in the resting state as well as in each task state. Most relationships with executive function were significantly different between the resting state and the task states, with tasks evoking new relationships that were not present in the resting state as well as strengthening one relationship that existed weakly in the resting state.

### Differences in functional brain organization between the resting state and executive function tasks

Effortful cognition, such as executive function, has long been thought to be facilitated by global integration across distributed brain regions (Chen & Deem, 2015; Dehaene et al., 1998; Kitzbichler et al., 2011). In accordance with this idea, it is well established that increased integration is related to better working memory and inhibition performance in adults (Bassett et al., 2009; Braun et al., 2015; Cohen et al., 2014; Cohen & D’Esposito, 2016; Meunier et al., 2010; Shine et al., 2016; Spielberg et al., 2015; Stanley et al., 2014). A small number of studies in children similarly demonstrate that integration is higher in working memory tasks relative to the resting state (Kolskår et al., 2018; Le et al., 2020), and that integration increases with increasing cognitive load in a task probing both working memory and response inhibition (Wang et al., 2020). Our findings build upon these prior studies in children and demonstrate increased whole brain and network-specific integration during both the SST and EN-back tasks compared to the resting state in late childhood. This indicates that the flexibility of network organization to adapt to executive function demands, well established in adults, is in place by late childhood.

It may be surprising that integration of the default mode network increased during tasks relative to the resting state, given the importance of default mode network anticorrelations with, or segregation from, other networks during effortful cognition, including response inhibition (Chung et al., 2020; Dwyer et al., 2014; Stevens et al., 2007) and working memory (Zhong et al., 2014). There is evidence that anticorrelations between the default mode and cognitive control networks increase with age (Barber et al., 2013; Baum et al., 2017; M. Chen et al., 2022; Keller et al., 2023; Stevens, 2016) and that task-based segregation of the default mode network may be less critical for executive function in children compared to adults (M. Chen et al., 2022). Therefore, it is possible that anticorrelations between the default mode network and task-relevant networks are not observed in children in this age range. Alternately, there is evidence for context-dependent alterations in connectivity between the default mode and cognitive control networks in adults (Cocchi et al., 2013; Fornito et al., 2012), such that stronger connectivity between the two systems may emerge during tasks (Fornito et al., 2012; Hearne et al., 2017) and can increase with greater complexity of task demands (Hearne et al., 2015, 2017). Our findings extend this notion into late childhood and could reflect both possibilities. For example, reconfiguration between the resting state and task states may not occur uniformly across the entire default mode network but may exhibit a combination of increases and decreases in connectivity based on the subsystem of the default mode network, the cognitive state, and neurodevelopmental trajectories. Future research should differentiate among the nuances of default mode network reconfiguration by cognitive demands across development.

### Tasks evoke unique relationships and strengthen existing relationships with executive function as compared to the resting-state

To understand how resting state and task-based functional connectivity differ in their relationships with executive function task performance, we evaluated brain-behavior relationships and tested for differences between the resting state and each task state. Prior studies using connectome-based predictive modeling have asserted that task-based functional connectivity evokes stronger relationships with behavior than resting state functional connectivity in adults (Greene et al., 2018, 2020; Jiang et al., 2020) and children (J. Chen et al., 2022; Zhao et al., 2023). We build upon this work to differentiate brain-behavior relationships that are established intrinsically from those that are evoked during the execution of tasks. We identified three types of relationships: those that are unique to the task states, those that exist weakly during the resting state and are strengthened during the task state, and those that may be unique to the resting state but for which further examination is warranted. We describe each relationship in detail below.

#### Executive function relationships unique to task-evoked functional connectivity

We most frequently observed that brain organization during a task evoked unique relationships with behavior that were not observed during the resting state. First, higher cingulo-parietal network dissociation index was significantly related to better performance on both the EN-back and SST tasks, yet no relationship between the cingulo-parietal network and behavior was seen during the resting state. Importantly, the relationships between executive function and cingulo-parietal network dissociation index were significantly stronger during both tasks compared to the resting state. The cingulo-parietal network, which includes regions in the posterior cingulate and posterior medial parietal cortex, lies in an integration zone between the default mode and fronto-parietal networks (Gordon et al., 2016; Leech & Sharp, 2014; Power et al., 2011; Yeo et al., 2011). A key region of the posterior cingulate cortex in the cingulo-parietal network is proposed to mediate shifts between internally- and externally-directed attention via changes in connectivity patterns with the default mode, fronto-parietal, and dorsal attention networks (Leech & Sharp, 2014). In accordance with this purported function, our results demonstrate that the cingulo-parietal network shifts to a more integrated state during executive function tasks compared to the resting state, and that greater integration is related to better performance on both the EN-back task and the SST, which is consistent with a successful shift in the locus of attention from internal to external. Our results additionally indicate that the degree of cingulo-parietal integration during the resting state, when the locus of attention is unconstrained, is not related to executive function task performance.

A similar relationship was observed for the cingulo-opercular network. Specifically, cingulo-opercular network dissociation index was significantly related to 2-back accuracy during the EN-back task, yet no relationship between the cingulo-opercular network and 2-back accuracy was observed during the resting state. Further, the relationship between 2-back accuracy and cingulo-opercular network dissociation index was significantly stronger during the EN-back task compared to the resting state. This finding accords with the conceptualization of the cingulo-opercular network as sustaining task control (Gratton, Sun, et al., 2018) and facilitating an “action mode” state generally oriented toward external processes (Dosenbach et al., 2024). These cingulo-parietal and cingulo-opercular relationships provide evidence that tasks evoke unique relationships with executive function that reflect on-line cognitive processes and cannot be detected during the resting state.

#### Executive function relationships strengthened during task-evoked functional connectivity

We identified one instance in which task-based functional connectivity strengthened a weak relationship with behavior present during the resting state. That is, lower dorsal attention network dissociation index during the EN-back task was related to higher 2-back accuracy, but lower dorsal attention network dissociation index during the resting state was only related to higher 2-back accuracy during the replication sample and not the discovery sample. Across both samples, the relationship between 2-back accuracy and dorsal attention network dissociation index was significantly stronger during the EN-back task compared to the resting state. This suggests that although the relationship to working memory performance during the resting state exists in some samples, this relationship is more consistent and stronger during task execution. During working memory, sustained representations of attentional priorities (Nee et al., 2013) and encoding of varying levels of load and task difficulty (Majerus et al., 2018) are facilitated by the dorsal attention network. Our results could indicate that less integration of the dorsal attention network with other networks facilitates these working memory-related processes. This relationship provides support for the notion that tasks may strengthen relationships that exist weakly during the resting state (Greene et al., 2018).

#### Executive function relationships that may be unique to the resting state

We found one instance in which resting state functional connectivity may exhibit a unique relationship with executive function, but the evidence is not conclusive. Specifically, lower fronto-parietal network dissociation index during the resting state was related to higher 2-back accuracy, but fronto-parietal network dissociation index EN-back task was not related to 2-back accuracy. This relationship between 2-back accuracy and fronto-parietal network dissociation index was only significantly different between the resting state and the EN-back task for the discovery sample, but not for the replication sample. Thus, we cannot conclusively state that this relationship is unique to the resting state. Working memory is a key skill engaged nearly constantly in everyday life and the importance of fronto-parietal engagement for working memory is well characterized (Barnes et al., 2016; Casey et al., 2018; Cohen et al., 2014; Engelhardt et al., 2019; Kharitonova et al., 2015; Kolskår et al., 2018; Murphy et al., 2020; Rosenberg et al., 2020). Resting state functional connectivity recapitulates patterns of frequent historical co-activation (Dosenbach et al., 2007; Gabard-Durnam et al., 2016; Kelly et al., 2008), so it is reasonable to speculate that greater expertise with working memory would instantiate stronger integration of the working memory-related fronto-parietal network during the resting state. The lack of an observed fronto-parietal relationship with 2-back accuracy during the EN-back task could perhaps result from the specific task design. The EN-back task included an affective component: on half of the 2-back blocks stimuli were emotional faces, while on the other half they were non-emotional places. It is possible that variability in fronto-parietal connectivity with affective regions between emotional and non-emotional task blocks precluded the ability to observe a consistent relationship with behavior during this task. In accordance with this idea, prior work has found that fronto-parietal functional connectivity with affective processing-related regions varies across emotion regulation strategies (Li et al., 2021). Further research is needed to understand how fronto-parietal integration contributes to working memory across the resting state and task contexts in late childhood.

## Conclusion

In summary, in a sample of children aged 9-10 years in the ABCD Study, we observed that relative to the resting state, functional brain network integration increased and segregation decreased during two tasks probing executive function, a stop signal task probing response inhibition and an emotional n-back task probing working memory. Additionally, we found that resting state and task-based network organization exhibited unique relationships with behavioral measures of response inhibition and working memory, which indicates unique rolls for intrinsic and task-online network organization in executive functions. Overall, this work shows that brain network organization is fine-tuned in the service of successful execution of tasks probing executive function in children. These task-related shifts in brain network organization both evoke new behaviorally-relevant brain signatures that are not observed during the resting state and strengthen relationships that exist weakly during the resting state.

The discovery that resting state and task-based functional connectivity exhibit unique relationships with behavior in late childhood has important implications. For example, the identification of executive function-related network features during the resting state may reflect that performance scales alongside a continuum of neurodevelopment from immature to mature, whereas executive function-related network features during task execution may reflect that performance scales with variance in engagement or strategy during on-line execution of a cognitive process. As there are likely strong reciprocal relationships between resting state and task-based functional connectivity, further attention should be paid to identifying how common and unique features that support executive functions shift based on cognitive demands. Both modalities offer complementary information toward a comprehensive understanding of the neurobiological mechanisms that give rise to complex behavior.

## Materials and Methods

### Participants

Behavioral and neuroimaging data from the baseline collection of the Adolescent Brain and Cognitive Development (ABCD) Study (Casey et al., 2018) was used for this project (Release 3.0; doi: 10.15154/1520591). The ABCD Study baseline collection consists of a demographically diverse sample of 11,875 children aged 9-10 years old. This data collection was conducted across 21 sites in the United States and is made available through the National Data Archive (NDA) at the National Institutes of Mental Health (NIMH). Neuroimaging data used in this study came from the ABCD-BIDS Collection 3165 v1.1.1 (https://collection3165.readthedocs.io/en/stable/) (Feczko, Earl, et al., 2021). Participants were excluded from the study if they did not have any functional MRI (fMRI) data in Collection 3165 (n = 1,885), their resting state fMRI data did not exist (n = 95) or were flagged for exclusion based on ABCD quality control criteria in the ABCD Release 3.0 notes (n = 1,911), or both SST and EN-back task data were flagged for exclusion (n = 1,488). Exclusion of SST data included both ABCD fMRI quality control criteria and criteria set by Garavan and colleagues (2020) to remove SST task errors. Exclusion of EN-back data included both ABCD fMRI quality control criteria and missing event files. Additionally, participants were excluded if they did not have sufficient timepoints remaining in both the resting state and at least one executive function task after removing high-motion timepoints during processing (n = 1,429). To provide matching discovery and replication datasets for this study, the final sample was divided into groups using the ABCD Reproducible Matched Samples (ARMS) designation provided in the Collection 3165 documentation (https://osf.io/7xn4f/; Feczko et al., 2020). Only matched groups 1 (*N* = 2,470) and 2 (*N* = 2,479) were used for this study. Therefore, the final sample consisted of *N* = 4,949 participants. See **Table 1** for demographic information for each sample.

### Measures

The ABCD neuroimaging protocol included a resting state scan and three cognitive task scans. The full ABCD study neuroimaging protocol, including descriptions of the fMRI tasks, is described in detail elsewhere (Casey et al., 2018). Relevant information for the current analyses is included here. All neuroimaging data, as well as task performance and demographic measures, were downloaded from the ABCD Study Release 3.0 on the NDA.

### Resting state

Up to four runs of resting state fMRI scans (5 minutes each) were collected from each subject, with a range of 12-20 minutes. During the resting state scans, subjects passively viewed a cross hair.

### Stop signal task (SST)

The SST probed response inhibition (Casey et al., 2018; Logan, 1994) (**Figure 4a**). In the task, subjects were instructed to press buttons to indicate if an arrow (the ‘go’ signal) was pointing left or right. On 16.67% trials, the go signal was interrupted with a vertical arrow (the ‘stop’ signal). The task utilized a staircase procedure that adjusted the time between the presentation of the go and stop signals, termed stop signal delay (SSD), so that over the course of the task run there were approximately 50% successful and 50% unsuccessful stop trials. The SSD was increased by 50ms following successful stop trials and decreased by 50ms following failed stop trials. Participants completed two runs of the SST, with each run containing 180 trials (150 go, 30 stop) and lasting a duration of 5 minutes and 49 seconds. Performance was indexed using the mean stop signal reaction time (SSRT), calculated as the mean reaction time on correct go trials minus the mean SSD (Logan et al., 1984). Average SSRT across both runs was similar across the discovery and replication samples (discovery: M = 292.42, SD = 62.25, range = 71.40 – 556.91; replication: M = 292.88, SD = 59.27, range = 32.20 – 482.96).

**Figure 4.**
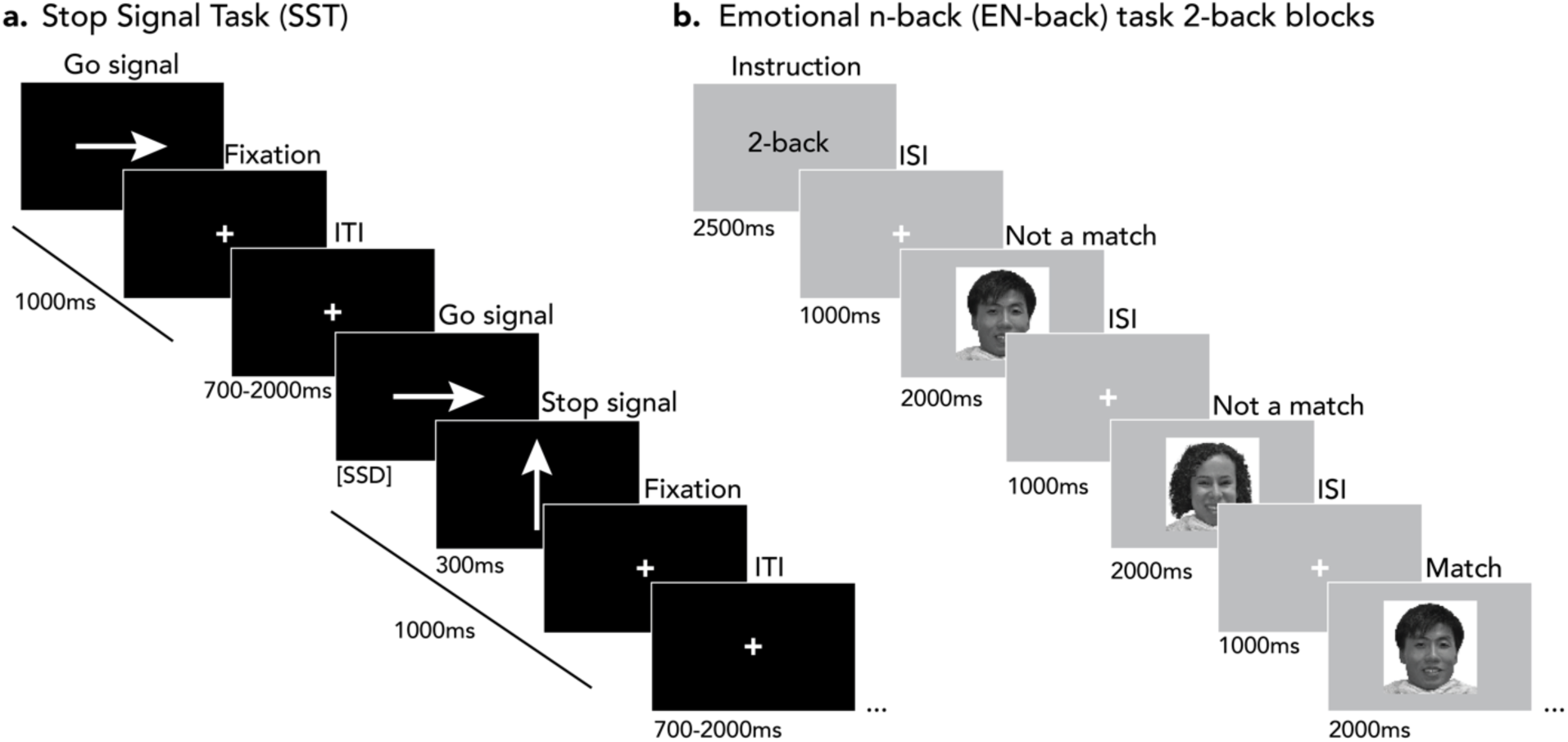
The fMRI tasks probing executive function. **(a)** The stop signal task (SST). Each trial (stimulus + fixation) was 1000ms and followed by an inter-trial interval (ITI) lasting between 700 and 2000ms. Go signals were terminated by the button press and the following fixation lasted the remainder of the 1000ms period. When go signals were followed by stop signals (300ms), the duration of the go signal was the ‘stop signal delay’ (SSD) determined by the staircase procedure. **(b)** 2-back blocks of the emotional n-back (EN-back) task. Each block began with a 2500ms instruction cue and then included a series of 2000ms stimuli each followed by a 1000ms inter-stimulus interval (ISI). Happy faces are shown as an example, but the task also included blocks of fearful faces, neutral faces, and places. This figure is an adaptation of the figures in (Casey et al., 2018).

### Emotional N-back (EN-back) task

The EN-back task probed working memory with emotional (i.e., faces) and non-emotional (i.e., places) stimuli (Barch et al., 2013; Casey et al., 2018) (**Figure 4b**). The task was presented in blocks, with one stimulus type per block. A cue at the beginning of each block (2500ms) indicated whether the task was 0-back or 2-back for that block. Participants viewed one stimulus at a time. During 0-back blocks, subjects pressed a button to indicate if each stimulus matched a target stimulus presented at the beginning of the block. During 2-back blocks, subjects pressed a button to indicate if each stimulus matched the stimulus presented two trials prior. Participants completed two runs of the EN-back task, with each run containing four 2-back blocks (40 trials) and four 0-back blocks (40 trials). For this analysis, only 2-back blocks were used to capture brain network organization during working memory. Performance was indexed as average accuracy on the 2-back blocks across both runs and was similar across the discovery and replication samples (discovery: M = 0.80, SD = 0.08, range = 0.61 – 0.99; replication: M = 0.80, SD = 0.08, range = 0.62 – 1.00). The inclusion of emotional stimuli in this analysis should not hinder the ability to identify brain-behavior relationships of working memory given that the EN-back task robustly engages core brain networks implicated in working memory for all conditions (Casey et al., 2018).

### Demographic measures

Demographic information collected included age, sex, race/ethnicity, parental education, family income, neighborhood disadvantage index, and parental marital status. Family income was collected in categorical bins of varying sizes (e.g., “$5,000 through $11,999”, “$75,000 through $99,999”), and the bins were converted into numeric quantities by taking the natural log of each of the following values: the highest value of the lowest bin (i.e., “Less than $5,000”: $5,000), the lowest value from the highest bin (i.e., “$200,000 and greater”: $200,000), and the midpoint from every other bin (e.g., “$75,000 through $99,999”: $87,499.50). Highest parental education was captured categorically (e.g., “GED or equivalent”) and each category was converted into a numeric value representing years of education (e.g., “GED or equivalent”: 12 years). An index of neighborhood disadvantage was calculated based on each participant’s home address (Fan et al., 2021; Kind et al., 2014) and provided by the ABCD Study. To align with recent studies (Rakesh et al., 2021; Sripada et al., 2021), we replicated the methods of Sripada and colleagues to calculate a composite socioeconomic status score from family income, parental education in years, and neighborhood disadvantage index via a confirmatory factor analysis in the lavaan R package (Rosseel, 2012).

### Image acquisition

Neuroimaging data were collected on three 3T scanner platforms (Siemens Prisma, *n*=3,480; General Electric 750, *n*=994; and Philips, *n*=475) with multi-channel head coils at 21 sites in the United States. Participants completed high-resolution 3D T1-weighted and T2-weighted images, diffusion weighted images, and functional MRI images from the resting state and the three fMRI tasks. The fMRI tasks were administered last and their order was counterbalanced across participants. The whole brain functional data were collected with a T2-weighted multiband EPI sequence with a slice acceleration factor of 6 (60 slices, 90×90 matrix, TR = 800 ms, TE = 30 ms, field of view 216 x 216mm, voxel dimensions: 2.4 mm x 2.4 mm x 2.4 mm). Acquisition parameters for the T1-weighted, T2-weighted, and diffusion images were harmonized across the three scanner platforms and full details are described elsewhere (Casey et al., 2018).

### Image processing

Image processing steps were implemented within the ABCD-BIDS pipeline (https://github.com/ABCD-STUDY/abcd-hcp-pipeline; (Fair et al., 2020; Feczko, Conan, et al., 2021; Feczko, Earl, et al., 2021; Glasser et al., 2013)). Source data were first converted into the standardized Brain Imaging Data Structure (BIDS) format and then processed. Structural MRI (T1-weighted) preprocessing steps included brain extraction, denoising and bias field correction, anatomical segmentation and cortical surface reconstruction in Freesurfer, and surface and volume registration to a reference template in standard space.

Functional MRI preprocessing steps included distortion correction with spin echo field maps, intensity normalization, and registration to the T1 and then the reference template. Data were mapped into individualized grayordinate space (“surface space”), smoothed with a 2mm full-width-half-maximum kernel, and then underwent additional processing via the *DCANBOLDProcessing* pipeline. Following (Marek et al., 2022), the BOLD processing steps included temporal notch filtering between 0.31-0.43Hz to attenuate respiration artifacts in the timeseries motion estimates (Fair et al., 2020), spectral interpolation of frames with a filtered FD > 0.3mm, time series detrending and normalization, and nuisance regression with six motion parameters and mean time series for white matter, cerebrospinal fluid and global signal, as well as derivatives and squares for each. Band-pass filtering between 0.008 and 0.1 Hz was then applied using a 2nd order Butterworth filter (Hallquist et al., 2013). Then, all functional runs were concatenated within each scan type (i.e., resting state, SST, or EN-back) and high motion scrubbing was implemented to remove timepoints with a filtered FD > 0.2mm from the time series (Power et al., 2012). Processing details can be found elsewhere (Feczko, Conan, et al., 2021; Marek et al., 2022).

After motion scrubbing, each concatenated timeseries was trimmed to be the same length across all subjects (i.e., 5 minutes (240 timepoints) for the resting state and the SST; 2 minutes (96 timepoints) for the EN-back task 2-back blocks). See **Table 2** for the number of subjects with usable data from each task. The concatenated time series data for the resting state and each task state was parcellated using the Gordon cortical parcellation (Gordon et al., 2016), which consists of 333 cortical surface parcels across 14 functional networks.

**Table 2.**
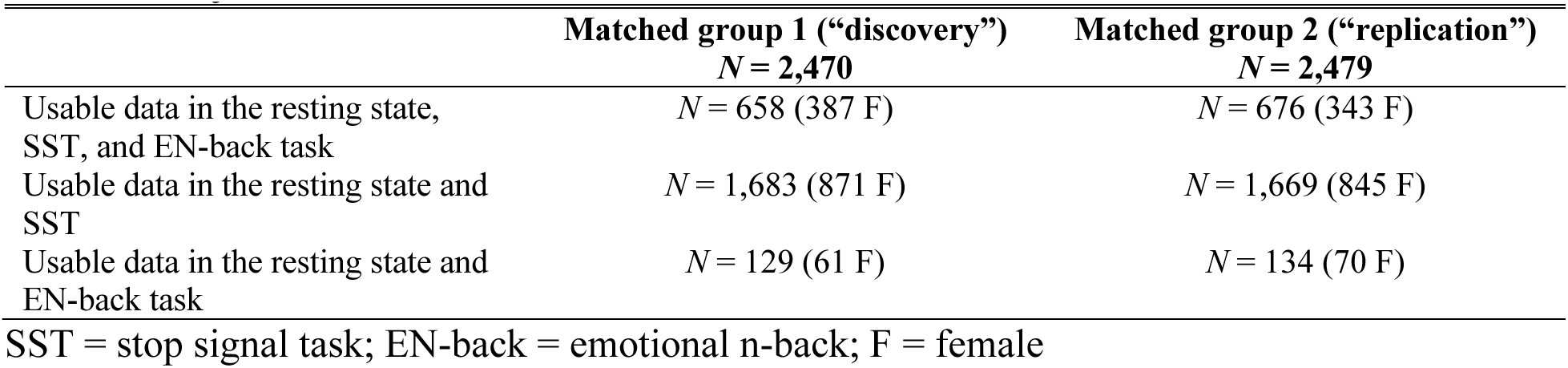
Subjects with data from each task state.

|  | Matched group 1 (“discovery”)<br>*N* = 2,470 | Matched group 2 (“replication”)<br>*N* = 2,479 |
| --- | --- | --- |
| Usable data in the resting state, SST, and EN-back task | *N* = 658 (387 F) | *N* = 676 (343 F) |
| Usable data in the resting state and SST | *N* = 1,683 (871 F) | *N* = 1,669 (845 F) |
| Usable data in the resting state and EN-back task | *N* = 129 (61 F) | *N* = 134 (70 F) |

SST = stop signal task; EN-back = emotional n-back; F = female

### Functional connectivity brain graph construction

Functional connectivity (FC) between each pair of parcels was estimated using Pearson’s correlation coefficient. This resulted in a separate 333 x 333 correlation matrix for each participant and each scan type (i.e., resting state, SST, and EN-back). Correlation matrices were Fisher z-transformed and thresholded to remove negative edges prior to the construction of brain graphs. Weighted, undirected brain graphs were calculated from the Fisher transformed, thresholded connectivity matrices using the igraph package in R (Csardi & Nepusz, 2006). Network assignments were pre-assigned from the Gordon parcellation (Gordon et al., 2016).

### Graph metric calculation

Graph metrics quantifying network segregation (modularity) and integration (global efficiency, node dissociation index) were calculated for each subject and each state. Modularity is a measure of the degree to which the brain segregates into distinct networks, with stronger within-network connections and weaker between-network connections (Bullmore & Bassett, 2011). A weighted variant of modularity was calculated using the pre-assigned network memberships from the Gordon parcellation (Gordon et al., 2016) with the igraph package in R (Csardi & Nepusz, 2006). Global efficiency is a measure of the ease of information transfer across the brain (without considering network membership). A weighted variant of global efficiency was calculated as the average of the shortest weighted path lengths defined on the inverse brain graph (Latora & Marchiori, 2001) and implemented with the brainGraph package in R (Watson, 2020).

Node dissociation index, which indexes network integration for specific nodes, was calculated for each node as the sum of weighted connections to nodes in every other network divided by the total weighted connections (Cary et al., 2016). Node dissociation index was then averaged across nodes in each network, which resulted in one value per network quantifying the integration of a network to other networks. Therefore, this metric will be referred to as ‘network dissociation index’. Finally, mean functional connectivity (FC) of the whole brain was calculated as the average of all edge values in the matrix remaining after removal of negative edges.

### Analyses

First, differences in functional brain network organization between the resting state and each task were quantified. Fourteen mixed effects models were estimated to test for differences in segregation (whole-brain modularity) and integration (whole-brain global efficiency, network dissociation index for each of 12 networks) between the resting state and both executive function task states. Mixed effects models with a random intercept of subject were implemented with the lme4 package in R (Bates et al., 2015). A Benjamini-Hochberg false discovery rate (FDR) correction at q < .05 (Benjamini & Hochberg, 1995) was applied across the 14 models separately for each resting state vs. task state comparison.

Then, for each task-evoked state, we tested if each brain metric during that state was related to task performance during that state (14 models per task). For the resting state, we tested if each brain metric was related to performance on both tasks (28 models). An FDR correction at q < .05 (Benjamini & Hochberg, 1995) was applied across the 14 models separately for each task performance measure (i.e., SSRT, 2-back accuracy) and each state (i.e., resting state, task). This resulted in four corrections with 14 p-values each.

Lastly, for all models in which we observed a significant relationship between a brain metric and task performance, we tested whether that relationship was significantly different between the resting state and the relevant task. Mixed effects models were estimated wherein a brain metric was predicted by an interaction between the scan type (rest, task) and task performance. A significant interaction term denoted a difference in the brain-behavior relationship between the resting state and the task-evoked state of interest. An FDR correction at q < .05 (Benjamini & Hochberg, 1995) was applied across all models.

In each model, age, sex, race/ethnicity, parental marital status, SES composite score, and scanner manufacturer were included as covariates. Mean FC was also included as a covariate, given the impact of differences in mean FC on network inferences (Hallquist & Hillary, 2019). All analyses were run separately and identically for both the discovery and replication samples.

## Acknowledgements

Data used in the preparation of this article were obtained from the Adolescent Brain Cognitive Development^SM^ (ABCD) Study (https://abcdstudy.org), held in the NIMH Data Archive (NDA). This is a multisite, longitudinal study designed to recruit more than 10,000 children aged 9-10 and follow them over 10 years into early adulthood. The ABCD Study® is supported by the National Institutes of Health and additional federal partners under award numbers U01DA041048, U01DA050989, U01DA051016, U01DA041022, U01DA051018, U01DA051037, U01DA050987, U01DA041174, U01DA041106, U01DA041117, U01DA041028, U01DA041134, U01DA050988, U01DA051039, U01DA041156, U01DA041025, U01DA041120, U01DA051038, U01DA041148, U01DA041093, U01DA041089, U24DA041123, U24DA041147. A full list of supporters is available at https://abcdstudy.org/federal-partners.html. A listing of participating sites and a complete listing of the study investigators can be found at https://abcdstudy.org/consortium_members/. ABCD consortium investigators designed and implemented the study and/or provided data but did not necessarily participate in the analysis or writing of this report. This manuscript reflects the views of the authors and may not reflect the opinions or views of the NIH or ABCD consortium investigators. The ABCD data repository grows and changes over time. The ABCD data used in this report came from <u>10.15154/1520591</u>. DOIs can be found at https://nda.nih.gov/abcd/abcd-annual-releases. The authors would also like to thank Tehila Nugiel for thoughtful feedback on the manuscript.

