## Supplementary Information for "Task-evoked functional connectivity exhibits novel and strengthened relationships with executive function relative to the resting state"

**Supplementary Table 1. Differences in brain organization between the resting state and the stop signal task (SST).**

| | Discovery sample | | | | | Replication sample | | | | |
|---|---|---|---|---|---|---|---|---|---|---|
| ***brain metric*** | ***b*** | ***st. b*** | ***SE*** | ***p*** | **adjusted-*p*** | ***b*** | ***st. b*** | ***SE*** | ***p*** | **adjusted-*p*** |
| Modularity | -0.04 | -1.12 | 0.0004 | <.001 | **<.001** | -0.04 | -1.10 | 0.0004 | <.001 | **<.001** |
| Global efficiency | 0.00 | 0.12 | 0.0001 | <.001 | **<.001** | 0.00 | 0.12 | 0.0001 | <.001 | **<.001** |
| Fronto-parietal NDI | 0.00 | -0.02 | 0.001 | 0.25 | 0.25 | 0.00 | -0.02 | 0.001 | 0.373 | 0.373 |
| Cingulo-opercular NDI | 0.04 | 0.69 | 0.001 | <.001 | **<.001** | 0.03 | 0.65 | 0.001 | <.001 | **<.001** |
| Salience NDI | 0.00 | 0.31 | 0.000 | <.001 | **<.001** | 0.00 | 0.34 | 0.0003 | <.001 | **<.001** |
| Cingulo-parietal NDI | 0.02 | 0.73 | 0.001 | <.001 | **<.001** | 0.02 | 0.74 | 0.001 | <.001 | **<.001** |
| Default mode NDI | 0.02 | 0.37 | 0.001 | <.001 | **<.001** | 0.02 | 0.39 | 0.001 | <.001 | **<.001** |
| Retrosplenial temporal NDI | 0.01 | 0.40 | 0.001 | <.001 | **<.001** | 0.01 | 0.42 | 0.001 | <.001 | **<.001** |
| Dorsal attention NDI | 0.03 | 0.69 | 0.001 | <.001 | **<.001** | 0.03 | 0.64 | 0.001 | <.001 | **<.001** |
| Ventral attention NDI | 0.03 | 0.91 | 0.001 | <.001 | **<.001** | 0.03 | 0.87 | 0.001 | <.001 | **<.001** |
| Somatomotor hand NDI | 0.02 | 0.46 | 0.001 | <.001 | **<.001** | 0.02 | 0.44 | 0.001 | <.001 | **<.001** |
| Somatomotor mouth NDI | 0.02 | 0.43 | 0.001 | <.001 | **<.001** | 0.01 | 0.38 | 0.001 | <.001 | **<.001** |
| Visual NDI | 0.17 | 1.45 | 0.002 | <.001 | **<.001** | 0.17 | 1.45 | 0.002 | <.001 | **<.001** |
| Auditory NDI | 0.02 | 0.65 | 0.001 | <.001 | **<.001** | 0.02 | 0.66 | 0.001 | <.001 | **<.001** |

NDI = network dissociation index; *b* = unstandardized beta; *st. b* = standardized beta; *SE* = standard error; bolded text denotes adjusted-*p* values < .05

**Supplementary Table 2. Differences in brain organization between the resting state and the emotional N-back (EN-back) task.**

| | Discovery sample | | | | | Replication sample | | | | |
|---|---|---|---|---|---|---|---|---|---|---|
| ***brain metric*** | ***b*** | ***st. b*** | ***SE*** | ***p*** | **adjusted-*p*** | ***b*** | ***st. b*** | ***SE*** | ***p*** | **adjusted-*p*** |
| Modularity | -0.09 | -2.64 | 0.00 | <.001 | **<.001** | -0.09 | -2.59 | 0.00 | <.001 | **<.001** |
| Global efficiency | 0.02 | 0.90 | 0.00 | <.001 | **<.001** | 0.02 | 0.90 | 0.00 | <.001 | **<.001** |
| Fronto-parietal NDI | 0.01 | 0.38 | 0.00 | <.001 | **<.001** | 0.01 | 0.45 | 0.00 | <.001 | **<.001** |
| Cingulo-opercular NDI | 0.12 | 2.29 | 0.00 | <.001 | **<.001** | 0.11 | 2.25 | 0.00 | <.001 | **<.001** |
| Salience NDI | 0.01 | 1.08 | 0.00 | <.001 | **<.001** | 0.01 | 1.13 | 0.00 | <.001 | **<.001** |
| Cingulo-parietal NDI | 0.05 | 1.55 | 0.00 | <.001 | **<.001** | 0.05 | 1.46 | 0.00 | <.001 | **<.001** |
| Default mode NDI | 0.12 | 2.41 | 0.00 | <.001 | **<.001** | 0.12 | 2.35 | 0.00 | <.001 | **<.001** |
| Retrosplenial temporal NDI | 0.02 | 0.83 | 0.00 | <.001 | **<.001** | 0.02 | 0.76 | 0.00 | <.001 | **<.001** |
| Dorsal attention NDI | 0.07 | 1.65 | 0.00 | <.001 | **<.001** | 0.07 | 1.66 | 0.00 | <.001 | **<.001** |
| Ventral attention NDI | 0.05 | 1.54 | 0.00 | <.001 | **<.001** | 0.05 | 1.49 | 0.00 | <.001 | **<.001** |
| Somatomotor hand NDI | 0.08 | 1.43 | 0.00 | <.001 | **<.001** | 0.07 | 1.37 | 0.00 | <.001 | **<.001** |
| Somatomotor mouth NDI | 0.04 | 1.02 | 0.00 | <.001 | **<.001** | 0.03 | 0.95 | 0.00 | <.001 | **<.001** |
| Visual NDI | 0.26 | 2.21 | 0.00 | <.001 | **<.001** | 0.25 | 2.16 | 0.00 | <.001 | **<.001** |
| Auditory NDI | 0.05 | 1.57 | 0.00 | <.001 | **<.001** | 0.05 | 1.58 | 0.00 | <.001 | **<.001** |

NDI = network dissociation index; *b* = unstandardized beta; *st. b* = standardized beta; *SE* = standard error; bolded text denotes adjusted-*p* values < .05

**Supplementary Table 3. Brain organization related to performance on the stop signal task (SST): SSRT.**

| | | Discovery sample | | | | | Replication sample | | | | |
|---|---|---|---|---|---|---|---|---|---|---|---|
| ***brain metric*** | ***cognitive state*** | ***b*** | ***st. b*** | ***SE*** | ***p*** | **adjusted-*p*** | ***b*** | ***st. b*** | ***SE*** | ***p*** | **adjusted-*p*** |
| Modularity | Resting state | -181.15 | -0.05 | 73.85 | 0.014 | 0.200 | -208.18 | -0.07 | 68.46 | 0.002 | **0.033** |
| Global efficiency | Resting state | 89.02 | 0.02 | 356.83 | 0.803 | 0.877 | 329.68 | 0.07 | 359.31 | 0.359 | 0.541 |
| Fronto-parietal NDI | Resting state | 74.44 | 0.04 | 44.78 | 0.097 | 0.338 | 46.10 | 0.02 | 43.39 | 0.288 | 0.541 |
| Cingulo-opercular NDI | Resting state | 76.99 | 0.05 | 36.50 | 0.035 | 0.220 | 74.18 | 0.05 | 35.18 | 0.035 | 0.145 |
| Salience NDI | Resting state | 84.39 | 0.02 | 114.93 | 0.463 | 0.810 | 81.79 | 0.02 | 109.34 | 0.455 | 0.568 |
| Cingulo-parietal NDI | Resting state | 90.48 | 0.04 | 45.56 | 0.047 | 0.220 | 42.89 | 0.02 | 42.21 | 0.310 | 0.541 |
| Default mode NDI | Resting state | 40.76 | 0.02 | 39.66 | 0.304 | 0.639 | 73.67 | 0.04 | 37.44 | 0.049 | 0.145 |
| Retrosplenial temporal NDI | Resting state | 26.63 | 0.01 | 54.16 | 0.623 | 0.872 | 10.47 | 0.00 | 52.83 | 0.843 | 0.843 |
| Dorsal attention NDI | Resting state | 22.52 | 0.01 | 36.67 | 0.539 | 0.839 | 22.52 | 0.01 | 34.01 | 0.508 | 0.568 |
| Ventral attention NDI | Resting state | 9.44 | 0.00 | 49.77 | 0.850 | 0.877 | 40.03 | 0.02 | 46.18 | 0.386 | 0.541 |
| Somatomotor hand NDI | Resting state | 4.26 | 0.00 | 27.62 | 0.877 | 0.877 | 24.65 | 0.02 | 26.10 | 0.345 | 0.541 |
| Somatomotor mouth NDI | Resting state | 14.64 | 0.01 | 39.62 | 0.712 | 0.877 | 24.50 | 0.01 | 38.76 | 0.527 | 0.568 |
| Visual NDI | Resting state | 20.58 | 0.02 | 19.61 | 0.294 | 0.639 | 36.85 | 0.05 | 18.93 | 0.052 | 0.145 |
| Auditory NDI | Resting state | 48.39 | 0.02 | 48.60 | 0.320 | 0.639 | 99.32 | 0.05 | 44.02 | 0.024 | 0.145 |
| Modularity | SST | 104.93 | 0.03 | 78.82 | 0.183 | 0.257 | 0.67 | 0.00 | 74.85 | 0.993 | 0.993 |
| Global efficiency | SST | 702.93 | 0.15 | 364.95 | 0.054 | 0.122 | 1426.37 | 0.31 | 338.42 | 0.000 | **0.000** |
| Fronto-parietal NDI | SST | 10.58 | 0.00 | 47.89 | 0.825 | 0.825 | 129.22 | 0.06 | 46.32 | 0.005 | **0.027** |
| Cingulo-opercular NDI | SST | 72.45 | 0.04 | 39.92 | 0.070 | 0.122 | 31.68 | 0.02 | 37.89 | 0.403 | 0.565 |
| Salience NDI | SST | -207.75 | -0.04 | 114.58 | 0.070 | 0.122 | 33.29 | 0.01 | 110.95 | 0.764 | 0.962 |
| Cingulo-parietal NDI | SST | -142.87 | -0.06 | 51.13 | 0.005 | **0.018** | -133.99 | -0.06 | 48.49 | 0.006 | **0.027** |
| Default mode NDI | SST | -131.33 | -0.08 | 38.17 | 0.001 | **0.004** | -44.80 | -0.03 | 36.28 | 0.217 | 0.355 |
| Retrosplenial temporal NDI | SST | -202.95 | -0.08 | 57.15 | 0.000 | **0.004** | -81.88 | -0.03 | 55.78 | 0.142 | 0.313 |
| Dorsal attention NDI | SST | -64.20 | -0.03 | 42.08 | 0.127 | 0.198 | 8.70 | 0.01 | 39.23 | 0.824 | 0.962 |
| Ventral attention NDI | SST | -102.72 | -0.04 | 54.59 | 0.060 | 0.122 | -61.26 | -0.03 | 50.82 | 0.228 | 0.355 |
| Somatomotor hand NDI | SST | 17.80 | 0.01 | 31.51 | 0.572 | 0.728 | 50.10 | 0.04 | 29.77 | 0.093 | 0.313 |

| | | | | | | | | | | | |
|---|---|---|---|---|---|---|---|---|---|---|---|
| Somatomotor mouth NDI | SST | 140.80 | 0.07 | 43.25 | 0.001 | **0.005** | 60.16 | 0.03 | 41.18 | 0.144 | 0.313 |
| Visual NDI | SST | 7.65 | 0.01 | 22.56 | 0.734 | 0.791 | -2.88 | 0.00 | 21.51 | 0.894 | 0.962 |
| Auditory NDI | SST | 17.33 | 0.01 | 44.54 | 0.697 | 0.791 | 61.17 | 0.03 | 43.16 | 0.157 | 0.313 |

NDI = network dissociation index; SSRT = stop signal response time; *b* = unstandardized beta; *st. b* = standardized beta; *SE* = standard error; bolded text denotes adjusted-*p* values < .05

**Supplementary Table 4. Brain organization related to performance on the Emotional N-back (EN-back) task: 2-back accuracy.**

| | | Discovery sample | | | | | Replication sample | | | | |
|---|---|---|---|---|---|---|---|---|---|---|---|
| ***brain metric*** | ***cognitive state*** | ***b*** | ***st. b*** | ***SE*** | ***p*** | **adjusted-*p*** | ***b*** | ***st. b*** | ***SE*** | ***p*** | **adjusted-*p*** |
| Modularity | Resting state | 0.16 | 0.04 | 0.09 | 0.072 | 0.202 | 0.15 | 0.04 | 0.09 | 0.083 | 0.246 |
| Global efficiency | Resting state | -0.14 | -0.02 | 0.44 | 0.749 | 0.982 | 0.09 | 0.01 | 0.46 | 0.854 | 0.854 |
| Fronto-parietal NDI | Resting state | -0.16 | -0.06 | 0.06 | 0.004 | **0.029** | -0.17 | -0.06 | 0.06 | 0.002 | **0.028** |
| Cingulo-opercular NDI | Resting state | 0.04 | 0.02 | 0.04 | 0.416 | 0.832 | -0.04 | -0.02 | 0.05 | 0.339 | 0.395 |
| Salience NDI | Resting state | 0.00 | 0.00 | 0.14 | 0.981 | 0.982 | 0.11 | 0.02 | 0.14 | 0.432 | 0.465 |
| Cingulo-parietal NDI | Resting state | -0.08 | -0.03 | 0.06 | 0.178 | 0.415 | -0.08 | -0.03 | 0.05 | 0.131 | 0.262 |
| Default mode NDI | Resting state | -0.09 | -0.04 | 0.05 | 0.072 | 0.202 | -0.08 | -0.03 | 0.05 | 0.088 | 0.246 |
| Retrosplenial temporal NDI | Resting state | 0.01 | 0.00 | 0.07 | 0.858 | 0.982 | 0.11 | 0.03 | 0.07 | 0.108 | 0.252 |
| Dorsal attention NDI | Resting state | -0.08 | -0.04 | 0.05 | 0.068 | 0.202 | -0.13 | -0.06 | 0.04 | 0.004 | **0.028** |
| Ventral attention NDI | Resting state | 0.04 | 0.01 | 0.06 | 0.544 | 0.951 | 0.07 | 0.02 | 0.06 | 0.214 | 0.323 |
| Somatomotor hand NDI | Resting state | 0.00 | 0.00 | 0.03 | 0.982 | 0.982 | -0.04 | -0.03 | 0.03 | 0.185 | 0.323 |
| Somatomotor mouth NDI | Resting state | 0.00 | 0.00 | 0.05 | 0.964 | 0.982 | -0.05 | -0.02 | 0.05 | 0.278 | 0.353 |
| Visual NDI | Resting state | 0.01 | 0.00 | 0.02 | 0.813 | 0.982 | 0.03 | 0.03 | 0.02 | 0.231 | 0.323 |
| Auditory NDI | Resting state | -0.19 | -0.06 | 0.06 | 0.001 | **0.019** | -0.10 | -0.03 | 0.06 | 0.085 | 0.246 |
| Modularity | EN-back task | -0.41 | -0.08 | 0.18 | 0.022 | 0.051 | -0.59 | -0.11 | 0.18 | 0.001 | **0.004** |
| Global efficiency | EN-back task | -2.60 | -0.40 | 0.70 | 0.000 | **0.001** | -1.33 | -0.21 | 0.68 | 0.053 | 0.124 |
| Fronto-parietal NDI | EN-back task | 0.03 | 0.01 | 0.09 | 0.723 | 0.779 | -0.09 | -0.03 | 0.09 | 0.344 | 0.438 |
| Cingulo-opercular NDI | EN-back task | 0.35 | 0.14 | 0.08 | 0.000 | **<.001** | 0.32 | 0.14 | 0.08 | 0.000 | **0.001** |
| Salience NDI | EN-back task | -0.27 | -0.04 | 0.25 | 0.277 | 0.485 | -0.21 | -0.03 | 0.24 | 0.386 | 0.446 |
| Cingulo-parietal NDI | EN-back task | 0.44 | 0.11 | 0.14 | 0.002 | **0.006** | 0.37 | 0.09 | 0.14 | 0.010 | **0.027** |
| Default mode NDI | EN-back task | 0.07 | 0.03 | 0.08 | 0.393 | 0.612 | 0.30 | 0.13 | 0.08 | 0.000 | **0.001** |
| Retrosplenial temporal NDI | EN-back task | -0.17 | -0.06 | 0.11 | 0.107 | 0.214 | -0.09 | -0.03 | 0.10 | 0.343 | 0.438 |
| Dorsal attention NDI | EN-back task | -0.37 | -0.16 | 0.08 | 0.000 | **<.001** | -0.29 | -0.14 | 0.07 | 0.000 | **0.001** |
| Ventral attention NDI | EN-back task | -0.08 | -0.02 | 0.13 | 0.562 | 0.656 | 0.02 | 0.00 | 0.14 | 0.907 | 0.907 |
| Somatomotor hand NDI | EN-back task | 0.20 | 0.10 | 0.07 | 0.003 | **0.008** | 0.10 | 0.05 | 0.07 | 0.129 | 0.226 |

| | | | | | | | | | | | |
|---|---|---|---|---|---|---|---|---|---|---|---|
| Somatomotor mouth NDI | EN-back task | -0.08 | -0.03 | 0.11 | 0.454 | 0.636 | 0.09 | 0.03 | 0.11 | 0.414 | 0.446 |
| Visual NDI | EN-back task | 0.00 | 0.00 | 0.06 | 0.961 | 0.961 | 0.10 | 0.06 | 0.06 | 0.094 | 0.188 |
| Auditory NDI | EN-back task | -0.06 | -0.02 | 0.10 | 0.528 | 0.656 | 0.13 | 0.05 | 0.09 | 0.146 | 0.227 |

NDI = network dissociation index; *b* = unstandardized beta; *st. b* = standardized beta; *SE* = standard error; bolded text denotes adjusted-*p* values < .05

**Supplementary Table 5. Cognitive state differences in relationships between brain organization and executive function task performance.**

| | | | Discovery sample | | | | | Replication sample | | | | |
|---|---|---|---|---|---|---|---|---|---|---|---|---|
| ***brain metric*** | ***cognitive state interaction term*** | ***task performance metric*** | ***b*** | ***st. b*** | ***SE*** | ***p*** | **adjusted-*p*** | ***b*** | ***st. b*** | ***SE*** | ***p*** | **adjusted-*p*** |
| CP NDI | SST vs. Resting state | SSRT | -0.0001 | -0.0018 | 0.00 | 0.000 | **<.001** | -0.000047 | -0.0015 | 0.00001 | 0.000 | **<.001** |
| FP NDI | EN-back vs. Resting state | 2-back accuracy | 0.03 | 0.89 | 0.01 | 0.044 | **0.044** | 0.00 | 0.11 | 0.01 | 0.803 | 0.803 |
| CO NDI | EN-back vs. Resting state | 2-back accuracy | 0.07 | 1.71 | 0.02 | 0.000 | **<.001** | 0.09 | 2.47 | 0.02 | 0.000 | **<.001** |
| CP NDI | EN-back vs. Resting state | 2-back accuracy | 0.04 | 1.36 | 0.01 | 0.000 | **<.001** | 0.04 | 1.30 | 0.01 | 0.000 | **<.001** |
| DA NDI | EN-back vs. Resting state | 2-back accuracy | -0.08 | -2.11 | 0.02 | 0.000 | **<.001** | -0.06 | -1.58 | 0.02 | 0.000 | **<.001** |

CP = cingulo-parietal; FP = fronto-parietal; CO = cingulo-opercular; DA = dorsal attention; NDI = network dissociation index; SST = stop signal task; EN-back = emotional n-back; SSRT = stop signal response time; *b* = unstandardized beta; *st. b* = standardized beta; *SE* = standard error; bolded text denotes adjusted-*p* values < .05
